# Persistent changes in nociceptor translatomes govern hyperalgesic priming in mouse models

**DOI:** 10.1101/2024.08.07.606891

**Authors:** Ishwarya Sankaranarayanan, Moeno Kume, Ayaan Mohammed, Juliet M Mwirigi, Nikhil Nageswar Inturi, Gordon Munro, Kenneth A Petersen, Diana Tavares-Ferreira, Theodore J Price

**Affiliations:** Pain Neurobiology Research Group, Department of Neuroscience, Center for Advanced Pain Studies, School of Behavioral and Brain Sciences, University of Texas at Dallas, Richardson, Texas; Hoba Therapeutics ApS, Copenhagen, Denmark

**Keywords:** chemotherapy-induced peripheral neuropathy, hyperalgesic priming, translating ribosome affinity purification, GPR88, Meteorin, IL6-mediated hyperalgesic priming

## Abstract

Hyperalgesic priming is a model system that has been widely used to understand plasticity in painful stimulus-detecting sensory neurons, called nociceptors. A key feature of this model system is that following priming, stimuli that do not normally cause hyperalgesia now readily provoke this state. We hypothesized that hyperalgesic priming occurs due to reorganization of translation of mRNA in nociceptors. To test this hypothesis, we used paclitaxel treatment as the priming stimulus and translating ribosome affinity purification (TRAP) to measure persistent changes in mRNA translation in Nav1.8+ nociceptors. TRAP sequencing revealed 161 genes with persistently altered mRNA translation in the primed state. We identified *Gpr88* as upregulated and *Metrn* as downregulated. We confirmed a functional role for these genes, wherein a GPR88 agonist causes pain only in primed mice and established hyperalgesic priming is reversed by Meteorin. Our work demonstrates that altered nociceptor translatomes are causative in producing hyperalgesic priming.

## Introduction

Nociceptors, the sensory neurons that innervate the entire bodies of animals and are responsible for detecting damaging or potentially damaging stimuli, display remarkable plasticity that is responsible for many pain states (*1–4*). For instance, in animal models and in human neuropathic pain patients, nociceptors can take on a state of hyperexcitability where they fire spontaneous action potentials with no apparent stimulus (*4–10*). This spontaneous activity is presumed to be the cause of the stabbing or shooting pain that chronic pain patients often experience. Injury to nociceptors causes changes in gene expression at the level of transcription and mRNA translation (*11, 12*). Transcriptional, translational, and post-translational signaling are important for regulation of the excitability of nociceptors (*2–4, 13*). Given the important role these neurons play in most forms of chronic pain (*5, 14, 15*), understanding nociceptor plasticity mechanisms can increase our understanding of why chronic pain occurs and can lead to opportunities for better treatment.

Injuries like surgical wounds, inflammatory mediators such as cytokines, and agents that can cause neuropathies, such as paclitaxel, provoke mechanical hyperalgesia and signs of ongoing pain in rodents and in humans. These pain-causing insults also produce a state referred to as hyperalgesic priming, wherein animals display a painful response to a dose of a stimulus, like prostaglandin E_2_ (PGE2), that does not normally cause pain in unprimed rodents (*16, 17*). The existence of this primed state, which can last from weeks to months after the initial insult, indicates that memory-like changes in the peripheral and/or central nervous system occur that maintain this primed state (*13, 16, 17*). Inhibitors of transcription and mRNA translation applied to the peripheral terminals and to the cell bodies of dorsal root ganglion (DRG) nociceptors can prevent development of hyperalgesic priming, and inhibitors of mRNA translation can reverse an established primed state (*18–29*). These findings suggest that the mRNA translational state of nociceptors is reorganized by the initial insult that causes the primed state and that this change continues even after the resolution of the initial pain state. However, this idea has not been formally tested.

We tested the hypothesis that hyperalgesic priming in mice is accompanied by a persistent change in the repertoire of translated mRNAs in nociceptors when mice are primed but showing no overt signs of mechanical hypersensitivity or ongoing pain. To do this, we used the translating ribosome affinity purification (TRAP) technique (*30, 31*). This technology uses a Cre-driven expression of the ribosomal L10a protein fused to enhanced green florescent protein (eGFP) to enable pulldown of translating ribosomes, coupled with RNA sequencing to assess the active translatome of a specific cell type (*30, 31*). We used the *Scn10a* gene as a Cre-driver because this gene is exclusively expressed in sensory neurons of mice and is greatly enriched in nociceptors versus other cell types (*32*). We have previously used this technique to gain insight into sex differences in translation in the DRG (*33*), difference in translatomes between the DRG and trigeminal ganglion (*34*), and to understand translational reorganization in nociceptors caused by the chemotherapeutic paclitaxel, but at the peak of paclitaxel-induced pain behaviors (*12*). Our findings support the hypothesis that translational reorganization in nociceptors is linked to the persistence of hyperalgesic priming. Moreover, we identified translationally up- and down-regulated mRNAs that can be activated to reveal the primed state, or can be increased to resolve the primed state, respectively. This work adds to a growing body of evidence that nociceptor plasticity is entrenched by multiple levels of gene expression regulation and requires active intervention to achieve resolution (*35–41*).

## Materials and Methods

### Animals

Rosa26^fsTRAP^ mice were crossed with Nav1.8^cre^ mice on a C57BL/6J genetic background to generate Nav1.8-TRAP mice, which express a fused eGFP-L10a protein in Nav1.8-positive neurons (*31–34*). Translating ribosome affinity purification (TRAP) experiments were conducted on female Nav1.8-TRAP littermates, while behavioral experiments were performed on both C57BL/6J mice and Nav1.8-TRAP mice that were 8-10 weeks old. The mice were group-housed with a maximum of 4 per cage and had food and water available ad libitum. They were maintained on a 12-hour light-dark cycle at a room temperature of 21 ± 2°C. All procedures were approved by the Institutional Animal Care and Use Committee at the University of Texas at Dallas.

### Injections

Paclitaxel (Sigma Aldrich, Y0000698) was dissolved in a 1:1 solvent mixture of Kolliphor EL (Sigma-Aldrich) and ethanol as previously described (*42, 43*). Female Nav1.8-TRAP mice were administered paclitaxel diluted in sterile Dulbecco’s phosphate buffered saline (DPBS; Thermo Scientific) or vehicle solvent intraperitoneally at a dosage of 4 mg/kg every other day, resulting in a cumulative dosage of 16 mg/kg. C57BL/6J mice were administered intraplantar injections of recombinant human IL-6 (R&D Systems, 206-IL) at a dosage of 0.1 ng or 100 ng prostaglandin E2 (PGE2; Cayman Chemical, cat# 14010) dissolved in sterile saline for priming experiments.

RTI-13951-33 hydrochloride (MedChem express), a GPR88 agonist, was dissolved in DMSO at a stock concentration of 5 mM, and then diluted in sterile saline and administered via intraplantar injection at doses of either 100 ng or 1 µg into the hind paw. Recombinant mouse Meteorin (rmMeteorin; R&D Systems, #3475) or vehicle control (DPBS) was administered once subcutaneously at 1.8 mg/kg using a 30-gauge needle. The investigator was blinded to the treatment during all injections.

### Behavioral Testing

Mice were habituated for 1 hour prior to testing for mechanical sensitivity in a clear acrylic behavioral chamber. Mechanical paw withdrawal thresholds were assessed using the Dixon up-down method with calibrated von Frey filaments (Stoelting), ranging from 0.16 g to 2 g, applied perpendicular to the mid-plantar surface of the hind paw (*44*). A positive response was defined as an immediate flicking or licking behavior upon application of the filament. The investigator was blinded to treatment conditions during all days of testing.

The Mouse Grimace Scale (MGS) was used to assess spontaneous pain-like behaviors in mice, as described by Langford et al. (*45*). Mice were scored in five facial categories: orbital tightening, nose bulge, cheek bulge, ear position, and whisker position. Each category was rated on a scale from 0 (not present), 1(moderately present) to 2 (severely present), and the average score was used to represent the overall level of facial changes. Mice were placed in an acrylic observation chamber, providing a clear view of the face for the observer. The investigator was blinded to the treatment conditions during the assessment. Grimace scores were assessed before testing for mechanical sensitivity to assure that grimacing responses were not a result of mechanical stimulation with von Frey filaments.

### Translating Ribosome Affinity Purification (TRAP)

Translating ribosome affinity purification was done as previously described by Tavares-Ferreira et.al. (*33*). Nav1.8-TRAP female mice (4 mice per condition) were euthanized by cervical dislocation and decapitation under isoflurane anesthesia, and their dorsal root ganglia (DRGs) were rapidly dissected in ice-cold dissection buffer (1× HBSS (Invitrogen, 14065006), 2.5 mM HEPES-NaOH [pH 7.4], 35 mM glucose, 5 mM MgCl2, 100 μg/ml cycloheximide, 0.2 mg/ml emetine). The DRGs were then transferred to ice-cold Precellys tissue homogenizing CKMix tubes containing lysis buffer (20 mM HEPES, 12 mM MgCl2, 150 mM KCl, 0.5 mM dithiothreitol (DTT), 100 μg/ml cycloheximide, 20 μg/ml emetine, 80 U/ml SUPERase•IN (Promega). Samples were homogenized using a Precellys® Minilys Tissue Homogenizer at 10-second intervals for a total of 80 seconds in a cold room (4°C). The lysate was then centrifuged at 2000 × g for 5 minutes to obtain the post nuclear fraction. Following this, 1% NP-40 and 30 mM 1,2-dihexanoyl-sn-glycero-3-phosphocholine were added, and the samples were centrifuged at 15,000 × g for 10 minutes to produce the post mitochondrial fraction. A 200 μl sample of this fraction was set aside for INPUT (bulk RNA-sequencing), while the remaining lysate was incubated with protein G-coated Dynabeads (Invitrogen) bound to 50 μg anti-GFP antibodies (HtzGFP-19F7 and HtzGFP-19C8, Memorial Sloan Kettering Centre) for 3 hours at 4°C with end-over-end mixing. The beads were washed with high salt buffer (20 mM HEPES, 5 mM MgCl2, 350 mM KCl, 1% NP-40, 0.5 mM DTT, and 100 μg/ml cycloheximide), and RNA was extracted using the Direct-zol kit (Zymo Research) following the manufacturer’s instructions. RNA yield was quantified using a Qubit (Invitrogen), and RNA quality was assessed using a Fragment Analyzer (Advanced Analytical Technologies).

Libraries were prepared according to the manufacturer’s instructions using the Total RNA Gold library preparation kit (with ribosomal RNA depletion) and sequenced on the NextSeq 2000 platform with 75-bp single-end reads. The sequencing was performed at the Genome Center, part of the University of Texas at Dallas Research Core Facilities.

### Data Analysis

RNA-seq read files (fastq files) were processed and analyzed following the methodology described by Tavares-Ferreira et al. (*33*). Quality assessment was conducted using FastQC (Babraham Bioinformatics). Reads were trimmed based on Phred scores and per-base sequence content. The trimmed reads were then aligned to the reference genome and transcriptome (Gencode vM16 and GRCm38.p5) using STAR v2.2.1. Relative abundances in Transcripts Per Million (TPM) for each gene in every sample were quantified using StringTie v1.3.5. Non-coding and mitochondrial genes were excluded from the analysis, and the TPMs for non-mitochondrial coding genes were re-normalized to sum to 1 million.

We selected 14,955 genes above the 30^th^ percentile in each INPUT sample, ensuring consistent detection of the transcriptome. Following this, we performed quantile normalization on all coding genes. For the IP samples, we calculated percentile ranks for the TPMs of these genes in each IP sample, focusing on genes above the 30^th^ percentile from the INPUT samples. From this set, 12,944 genes were identified as consistently detected in the translatome (IP samples), based on their expression at or above the 15^th^ percentile in each sample and were quantile normalized.

Differential gene expression analysis was performed by calculating the log2-fold change (based on median TPM values), strictly standardized mean difference (SSMD), and the Bhattacharya coefficient. This modified coefficient ranges from 0, indicating completely identical distributions, to +1 or -1, indicating entirely non-overlapping distributions, with the sign determined by the log-fold change value. TPMs for all genes are shown in **Supplementary File 1.**

### Statistics

Statistical analyses for behavioral data were conducted using GraphPad Prism 10. Data visualization was performed in Python (version 3.7, Anaconda distribution). Results are presented as mean ± SEM and were analyzed using one-way or two-way ANOVA followed by post hoc tests for between-group comparisons. Effect sizes were determined by subtracting behavior scores for each time point from baseline measures. Absolute values were summed up and plotted for each group. All data are represented as mean ± SEM with p<0.05 considered significant. Detailed statistical outcomes are provided in the figure legends. Bioinformatics analysis and data visualization coding were done in Python (version 3.7, Anaconda distribution). Schematic illustrations were created with BioRender (BioRender.com).

## Results

Our previous work demonstrated changes in nociceptor translatomes associated with paclitaxel neuropathy at the peak time point for mechanical hypersensitivity (*12*). We have also shown that after the resolution of paclitaxel-induced pain behaviors, mice demonstrate hyperalgesic priming (*46*) suggesting a persistent change in nociceptors that maintains this primed state (*13, 16, 17*). While there are many types of stimuli that can be used to induce hyperalgesic priming, the paclitaxel model is ideal for assessing the translatome of nociceptors because it is a systemic treatment that affects all DRGs, which need to be pooled to obtain enough material for TRAP-seq experiments (*12, 33*). To explore nociceptor-specific changes in mRNA translation after overt pain behaviors induced by paclitaxel have resolved, but when mice are still primed, we used paclitaxel-treated female Nav1.8-TRAP mice **(Figure 1A)**. Behavioral testing revealed significant mechanical hypersensitivity in paclitaxel-treated mice compared to vehicle controls, which persisted for nearly 50 days before returning to baseline **(Figure 1B)**. DRGs were dissected from paclitaxel- and vehicle-treated animals at day 59, followed by immunoprecipitation of translating ribosomes from DRG neurons, mRNA purification, and RNA-seq. We acquired both the Input sample (IN sample), which corresponds to the total mRNA present in the DRG, and the IP sample, which corresponds with the mRNA bound to eGFP-tagged ribosomes that are actively being translated **(Figure 1C)**.

**Figure 1:**
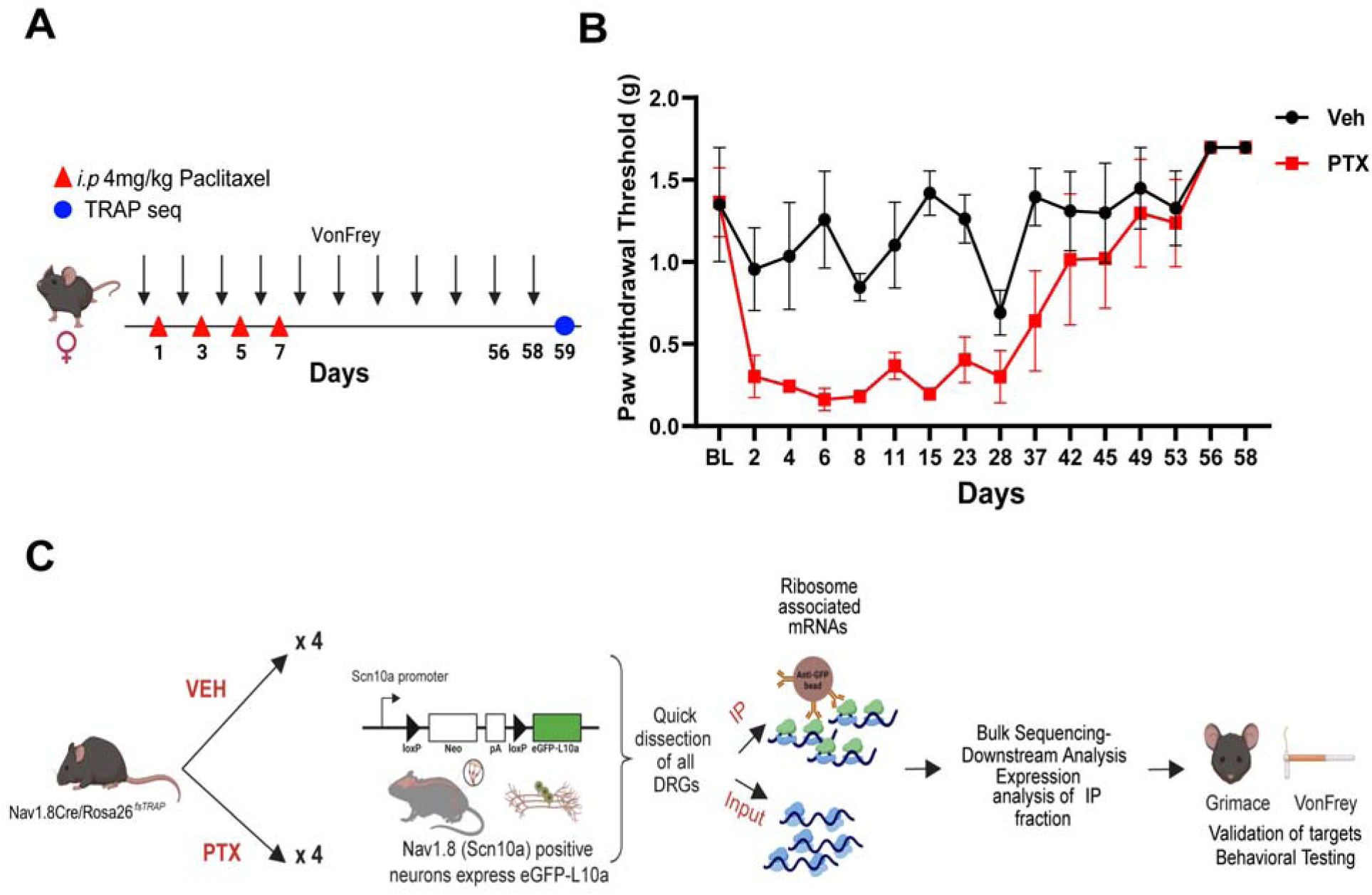
Cell type-specific gene expression during the resolution phase of paclitaxel-induced mechanical hypersensitivity. **A)** Schematic representation of the dosing paradigm for paclitaxel treatment and behavioral testing using von Frey filaments. **B)** Paw withdrawal thresholds were reduced with the administration of paclitaxel in a cohort of female Nav1.8cre/Rosa26^fsTRAP^ mice. **C)** Outline of the workflow for TRAP sequencing. Nav1.8cre/Rosa26^fsTRAP^ mice were administered vehicle or paclitaxel, and we euthanized on day 59 day upon returning to baseline mechanical thresholds. Whole DRGs were dissected and homogenized, and total RNA (INPUT) was extracted from the lysate. mRNAs bound to ribosomes (IP) were further isolated using eGFP-coated beads. Samples were sequenced and potential genes were validated using behavioral testing.

We conducted hierarchical clustering and heatmap analysis using the correlation coefficients of coding gene TPMs. This analysis revealed that the IN and IP samples were segregated into two distinct molecular profiles, demonstrating a clear separation between the actively translated mRNAs captured by TRAP-seq and the bulk RNA population analyzed by RNA-seq **(Figure 2A)**. We also identified a clear separation of each biological replicate between the TRAP-seq and bulk RNA-seq from the heatmap analysis **(Figure 2B)**. We identified high correlation coefficients across gene TPMs between input and IP samples (shown), revealing strong reproducibility across experiments **(Figure 2C)**. Percentile ranks were calculated for gene expression levels for each RNA-seq and TRAP-seq sample. Using these order statistics, we identified 14955 genes (>= 30^th^ percentile) that were consistently detected in at least one of the RNA-seq samples, and for the TRAP-seq samples, we detected 12944 genes (>= 15^th^ percentile). These findings align with previous mouse DRG RNA-seq and TRAP-seq studies (*12, 33, 34*). We plotted the empirical probability densities of coding gene TPMs and observed a bimodal distribution, distinguishing genes that are consistently detected from those that are lowly expressed. To account for differences in sequencing depth and ensure sample comparability, we applied quantile normalization to the TPM expression levels, represented as qnTPMs **(Figure 2D).** To verify the specificity of TRAP-seq for purifying translating mRNAs from nociceptors, we examined sets of control genes known to be enriched in either neuronal or non-neuronal cells within the DRG. Nociceptor mRNAs, such as *Calca, Trpv1, Scn10a,* and *Prph*, exhibited higher relative abundance in the IP sample. Conversely, non-neuronal genes, including glial markers like *Mpz, Mbp*, and *Gfap*, were found to be de-enriched in the IP samples **(Figure 2E)**. We identified differentially expressed (DE) genes by calculating the log2-fold change across the samples. Additionally, we employed two other statistics: the strictly standardized mean difference (SSMD) (*47, 48*) of TPM percentile ranks and the Bhattacharyya coefficient (BC) of qnTPMs. These measures quantified the effect size and controlled for within-group variability. For a gene to be considered differentially expressed, we applied stringent criteria: | log2-fold change | > 0.58 (equivalent to a fold change > 1.5), | SSMD | > 0.97, and BC < 0.5. We then plotted the SSMD values against the log2-transformed fold change for autosomal genes in IP samples **(Figure 2F)**.

**Figure 2:**
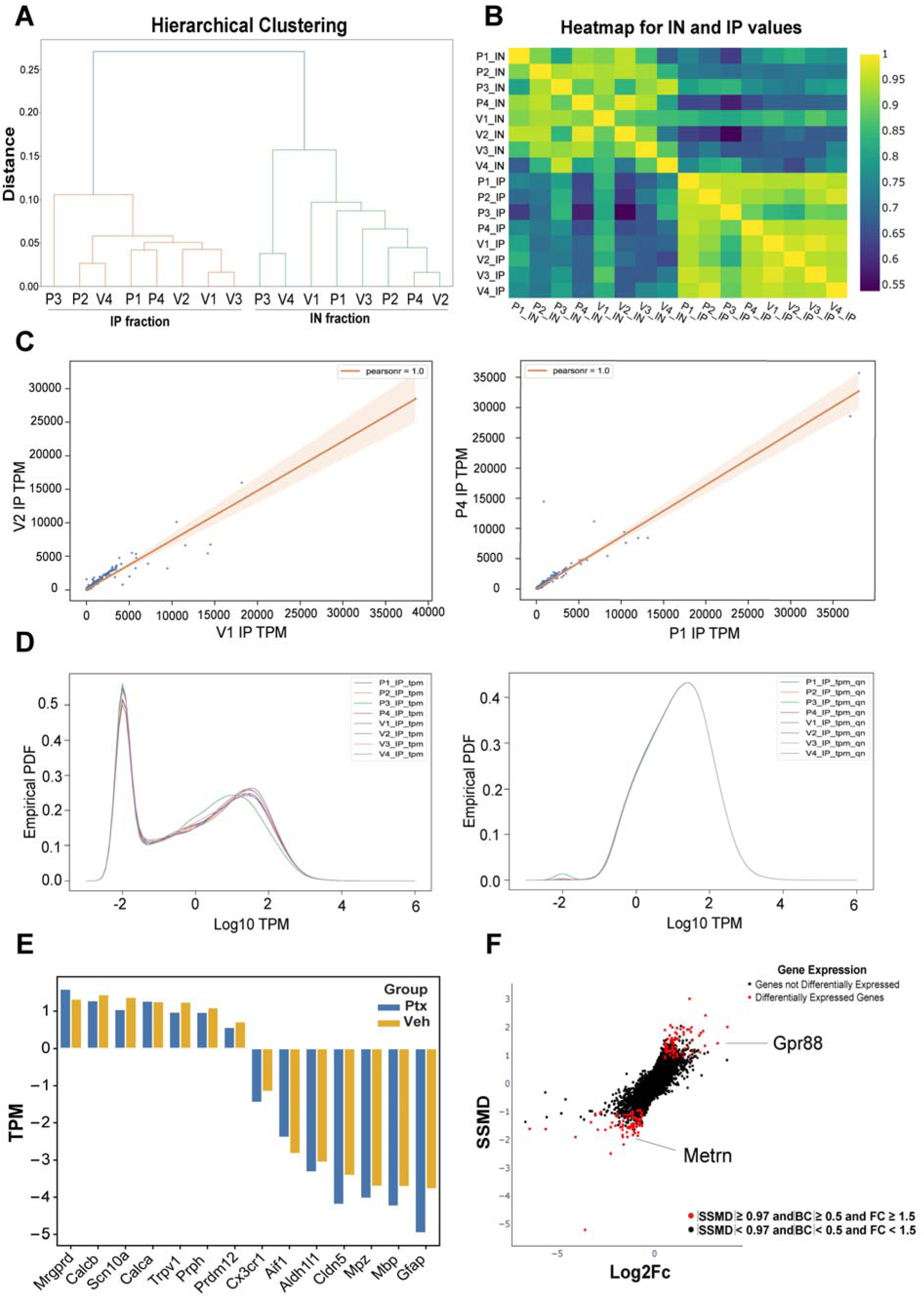
Nociceptor-specific translating ribosome affinity purification (TRAP) sequencing after resolution of paclitaxel-induced mechanical hypersensitivity: **A)** Hierarchical clustering using the correlation coefficients showing separation between the IN and the IP samples. **B)** Heatmap plot revealing a clear distinction between the biological replicates between the TRAP-seq and the INPUT samples. **C)** Linear correlation plots show a higher correlation between the IP samples. **D)** Empirically probability distribution of all coding genes using the raw TPMs and quantile normalized TPMs of all IP samples. **E)** Higher expression is observed of neuronal genes (*Mrgprd, Calcb, Scn10a, Calca, Trpv1, Prph*) compared to non-neuronal genes (*Prpdm12, Cx3cr1, Aif1, Mpz, Mbp, Gfap*) showing neuron-specific enrichment using TRAP. **F)** Differentially translated mRNAs in the IP sample are shown by a dual-lightened plot against SSMD and log2-fold change.

We identified 122 differentially expressed genes in the IN samples, with *Dusp14, Eef2kmt*, and *Mrgprx1* upregulated and *Mrgprh* downregulated. In the mRNAs associated with translating ribosomes (IP samples), we found a total of 160 differentially expressed genes, with 79 upregulated and 81 downregulated **(Supplementary File 2)**. We observed only four genes that overlapped between the IP and IN samples with genes such as *Mok, Tmem254a, Tmem254b, and Nup37*. Additionally, we found that the genes *Mok, Prcc, Ibtk, Fhdc1, Cars*, and *Ctu2* overlapped between our previous study doing TRAP sequencing at the peak of paclitaxel-evoked pain and the resolution of paclitaxel-induced pain (*12*). *Mok* is part of the MAP kinase pathway, suggesting continued involvement of this post-translational pathway from the early pain state to the time point of the primed state. Among the upregulated genes in the IP samples, several were already implicated in the sensitization of nociceptors, such as the prostaglandin E receptor 1 (*Ptger1*) and transforming growth factor beta1 (*Tgfb1*) (*49–53*) (**Figure 3A**). Conversely, the downregulated genes included several involved in neuronal function, such as sodium voltage-gated channel alpha subunit 3 (*Scn3a*) and solute carrier family 1 member 2 (*Slc1a2*) (**Figure 3B**).

**Figure 3:**
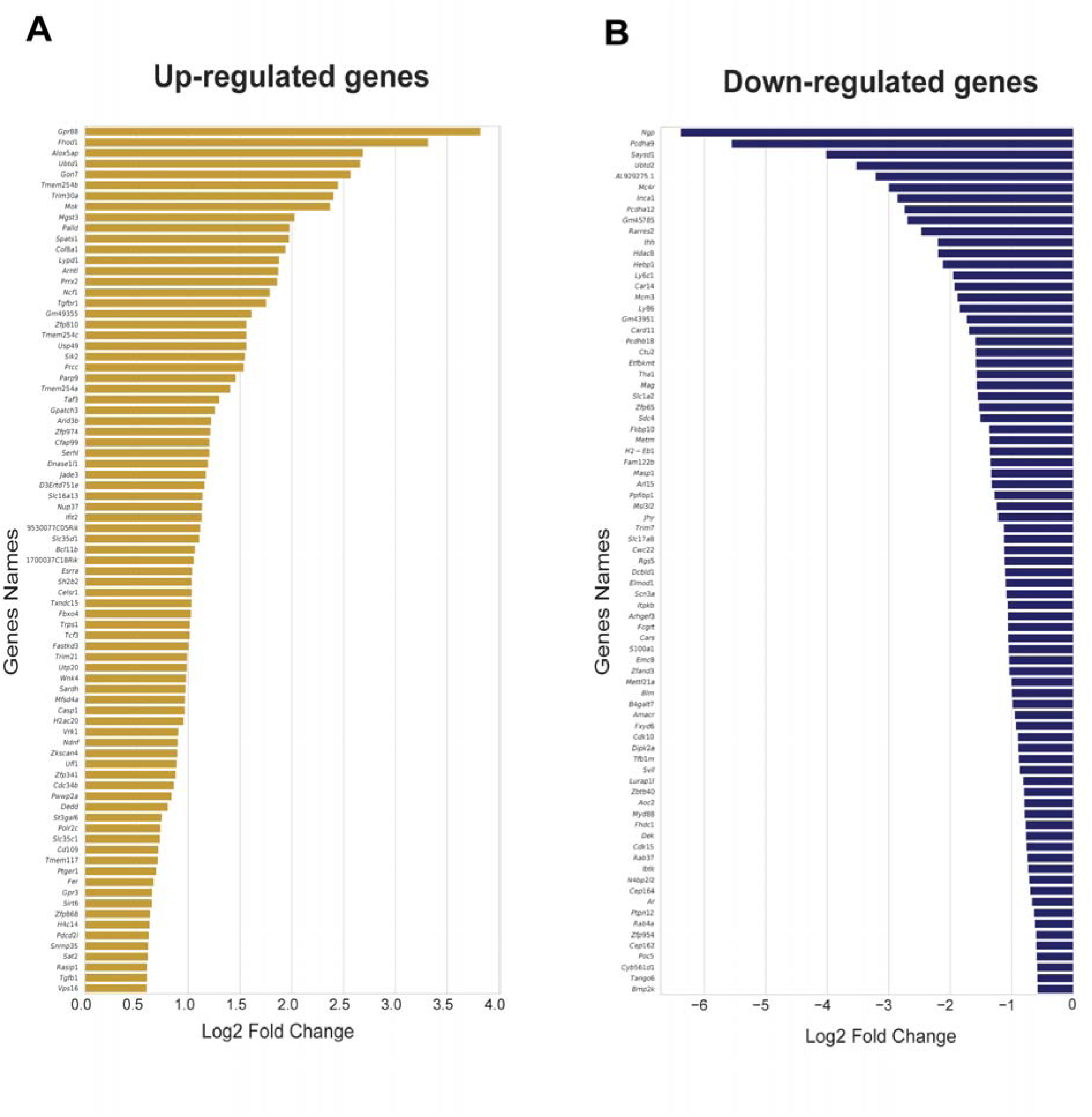
Differentially translated nociceptor-specific mRNAs after paclitaxel treatment. **A)** Bar plot showing a log2-fold change of all upregulated differentially translated mRNAs in the IP sample. **B)** Bar plot showing log 2-fold change of all downregulated genes in the IP samples.

We then focused on the most highly upregulated DE gene in the IP sample, *Gpr88* (**Figure 4A**), a gene encoding a G-protein-coupled receptor with no known endogenous ligand that is highly expressed in the striatum and cortex, associated with neurological disorders, and has several recently described synthetic agonists (*54*). We investigated whether a GPR88 agonist could be used in a similar manner to PGE2 to reveal a primed state by inducing mechanical hypersensitivity and grimacing only in primed animals. To reduce the time needed to assess the role of GPR88 in hyperalgesic priming and to test the generalizability of the target in other priming models, we used interleukin 6 (IL-6) as the priming stimulus. IL-6 is a well-studied pro-inflammatory mediator known to sensitize neurons in the DRG and produce an initial pain state that lasts for only about 5 days (*55–58*). We administered IL-6 into the paw and performed von Frey testing until the animals returned to baseline. Once the animals were back to baseline, they were injected with either a GPR88 agonist, RTI-13951-33 hydrochloride (100 ng or 1 µg), or 100 ng PGE2 **(Figure 4B)**. Intraplantar administration of IL-6 to female mice produced mechanical hypersensitivity which returned to baseline within a week post-administration. Mice previously treated with IL-6 and primed showed mechanical hypersensitivity when given an intraplantar injection of RTI-13951-33, but mice previously treated with vehicle and not primed did not respond to even the 1 µg dose of RTI-13951-33 **(Figure 4C)**. Primed mice also responded to the positive control 100 ng PGE2. Effect size calculation showed that only mice primed with IL-6 responded to either RTI-13951-33 or PGE2 **(Figure 4D)**. We also investigated grimacing behavior in these animals and observed that primed animals injected with either dose of the GPR88 agonist exhibited grimacing behaviors comparable to animals treated with IL-6 and PGE2 **(Figure 4E)**. This finding demonstrates that increased nociceptor translation of *Gpr88* is linked to a behavioral response to a GPR88 agonist only observed in primed mice.

**Figure 4:**
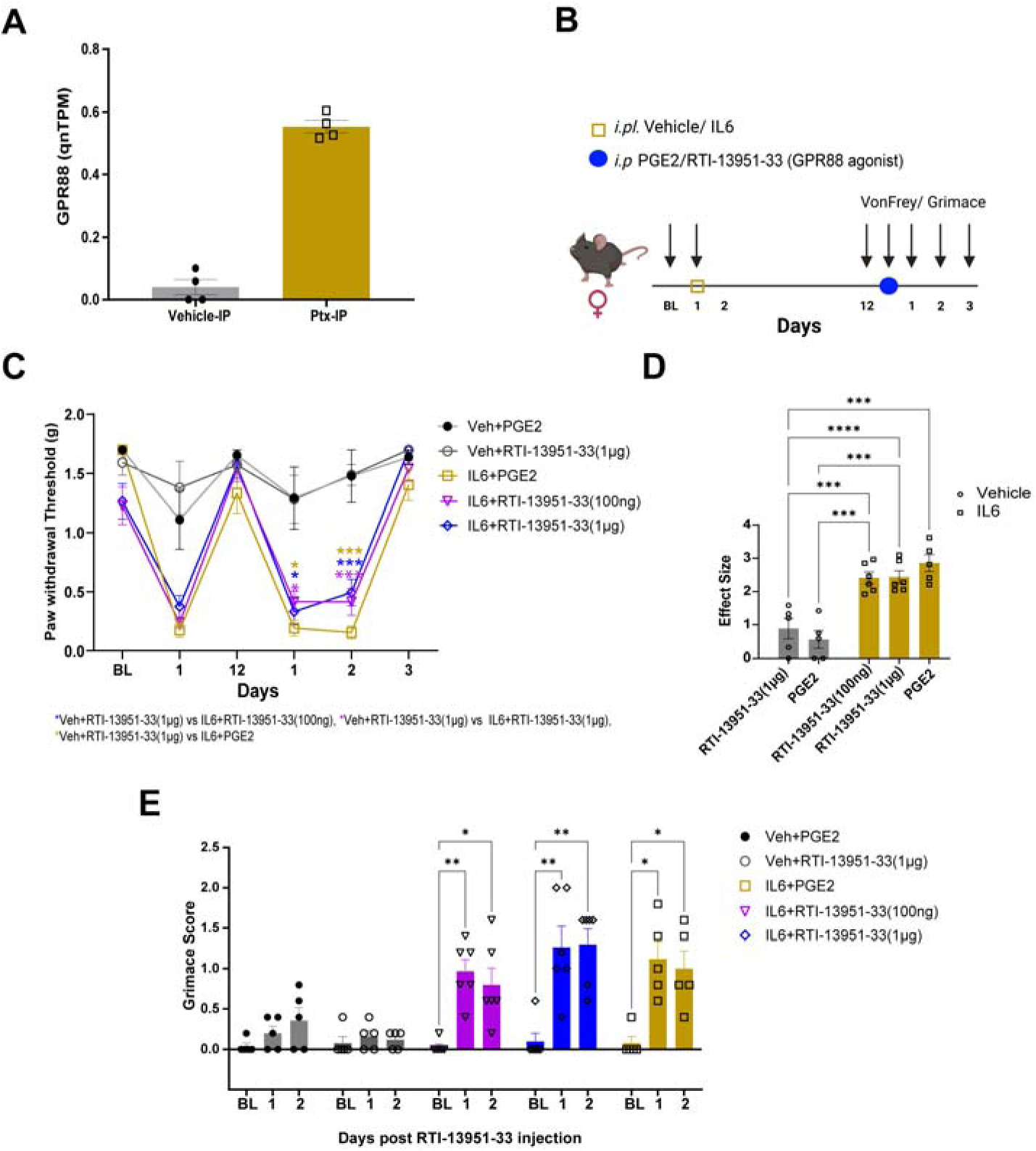
GPR88 causes mechanical hypersensitivity in primed animals after IL-6 treatment. **A)** Expression of *Grp88* mRNA is higher in paclitaxel-treated animals after the resolution of paclitaxel-induced pain behaviors. **B)** Schematic representation of dosing paradigm for intraplantar injection of IL-6, PGE2, or RTI-13951-33 (GPR88 agonist) and von Frey assessment. **C)** 100 ng and 1 µg RTI-13951-33 induce mechanical hypersensitivity only in primed mice that were previously treated with IL-6 (Two-way ANOVA, F= 5.173, p <0.0001, Tukey’s multiple comparison test, (Vehicle+PGE2, N=5, IL-6+1 µg RTI-13951-33, N=6, IL-6+100 ng RTI-13951-33, N=6, Vehicle+1 µg RTI-13951-33, N=5), Day 1 post PGE2 or RTI-13951-33, IL-6+PGE2 vs. Veh+1 µg RTI-13951-33, p-value = 0.0205, IL-6+1 µg RTI-13951-33 vs. Veh+1 µg RTI-13951-33, p-value = 0.0311, IL-6+100 ng RTI-13951-33 vs. Veh+1 µg RTI-13951-33, p-value = 0.0471, Day 2 post PGE2 or RTI-13951-33 administration IL-6+PGE2 vs. Veh+1 µg RTI-13951-33, p-value = < 0.0001, IL-6+1 µg RTI-13951-33 vs. Veh+1 µg RTI-13951-33, p-value = 0.0005, IL-6+100 ng RTI-13951-33 vs. Veh+1 µg RTI-13951-33, p-value = 0.0003, Veh+PGE2 vs. IL-6+PGE2, p-value = 0.0153, Veh+PGE2 vs. IL-6+1 µg RTI-13951-33, p-value = 0.0403, Veh+PGE2 vs. IL-6+100 ng RTI-13951-33, p-value = 0.0287. Data represents mean ± SEM. Significance represented as IL-6+1 µg RTI-13951-33 vs. Veh+1 µg RTI-13951-33 (pink*), IL-6+100 ng RTI-13951-33 vs. Veh+1 µg RTI-13951-33 (blue*), Veh+RTI-13951-33 vs. IL-6+PGE2 (yellow*). **D)** Effect size was determined by calculating the sum of the cumulative difference between the value for each time point post-GPR88 agonist administration and the day 12 baseline value. We observed a significant difference for both 100 ng and 1 µg RTI-13951-33 treatment between the Vehicle group and IL-6 primed animals. (Two-way ANOVA, p-value = <0.0001, Tukey’s multiple comparison test, 100 ng RTI-13951-33:IL-6 vs. 1 µg RTI-13951-33: Vehicle, p-value = 0.0002, 100 ng RTI-13951-33:IL-6 vs. PGE2:Vehicle, p-value = 0.0003, 1 µg RTI-13951-33:Vehicle vs. 1 µg RTI-13951-33:IL-6, p-value = <0.0001, 1 µg RTI-13951-33:Vehicle vs. PGE2:IL-6, p-value = 0.0002, 1 µg RTI-13951-33:IL-6 vs. PGE2:Vehicle, p-value = 0.0002, PGE2:Vehicle vs. PGE2:IL-6, p-value = <0.0001. **E)** Grimacing behavior was observed to be significantly different in animals treated with RTI-13951-33 compared to its baseline measurement. (Two-way ANOVA, F=4.834, p-value = 0.0003, Dunnett’s multiple comparison test, Day1 IL-6+PGE2 vs Baseline, p-value = 0.0261, IL-6+1 µg RTI-13951-33 vs Baseline, p-value = 0.0051, IL-6+100 ng RTI-13951-33 vs Baseline, p-value = 0.0022, Day2, IL-6+PGE2 vs Baseline, p-value = 0.0358, IL-6+1 µg RTI-13951-33 vs Baseline, p-value = 0.0024, IL-6+ 100 ng RTI-13951-33 vs Baseline, p-value = 0.0282). Data represents mean +/- SEM.

We identified the *Metrn* gene, which encodes Meteorin, a secreted protein with no known receptor originally reported to be expressed in neural progenitors and in astrocytes including radial glia (*59*), as one of the downregulated genes in the IP sample **(Figure 5A)**. Previous studies have demonstrated an antinociceptive effect of Meteorin in neuropathic pain models in mice and rats (*42, 60, 61*), but it has never been assessed in hyperalgesic priming models. We used the IL-6 priming model to assess whether recombinant mouse Meteorin (rmMTRN) treatment could reverse hyperalgesic priming. Once animals returned to baseline mechanical thresholds after IL-6 administration, a single subcutaneous injection of rmMTRN (1.8 mg/kg) was administered one hour before the intraplantar PGE2 injection **(Figure 5B)**. Treatment with rmMTRN led to significant alleviation of mechanical hypersensitivity caused by PGE2 treatment **(Figure 5C and D)**. rmMTRN treatment also alleviated PGE2-induced grimacing behavior **(Figure 5E)**.

**Figure 5:**
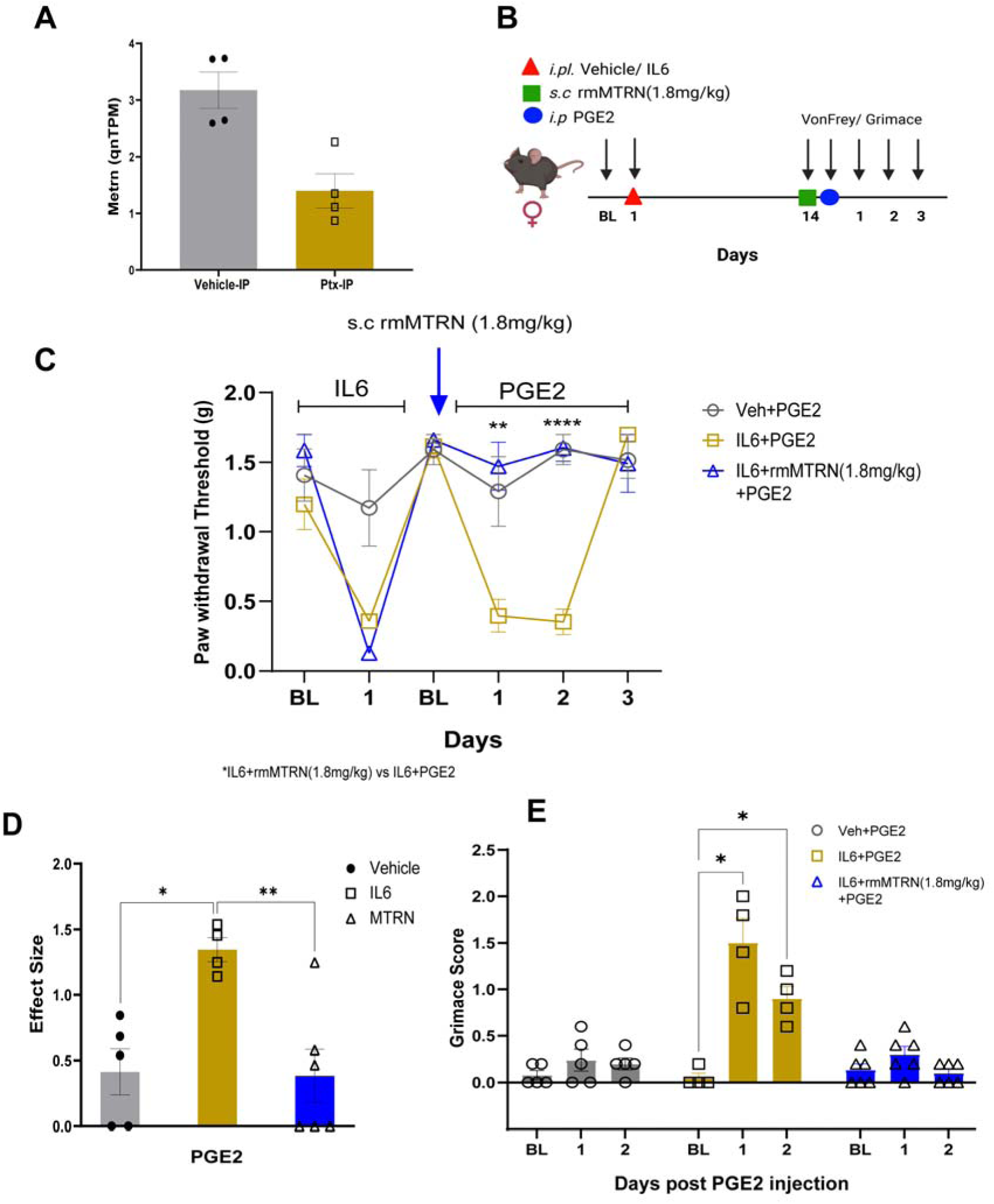
Meteorin reverses hyperalgesic priming in animals primed with IL-6. **A)** Quantile normalized expression of *Metrn* is observed to be decreased in paclitaxel-treated Nav1.8cre/Rosa26^fsTRAP^ mice after the resolution of paclitaxel-induced pain behaviors. **B)** Diagram showing the dosing paradigm of IL-6, rmMTRN, and PGE2 and behavioral testing schedule. **C)** 1.8mg/kg rmMTRN administration an hour prior to PGE2 injection attenuated the development of mechanical hypersensitivity in IL-6-primed animals. (Two-way ANOVA, F = 24.77, p-value = <0.0001, Tukey’s multiple comparison test, Day1 post PGE2 injection, IL-6+PGE2 vs. IL-6+rmMTRN (1.8 mg/kg)+PGE2, p-value = <0.0023, Day2 post PGE2 injection, IL-6+PGE2 vs. IL-6+rmMTRN (1.8 mg/kg)+PGE2, p-value = < 0.0001, IL-6+PGE2, N=4, Veh+PGE2, N=5, IL-6+rmMTRN (1.8 mg/kg)+PGE2, N=6). **D)** Effect size was calculated from the first administration of rmMTRN and was observed to be significant between primed animals treated with PGE2 and primed animals administered with rmMTRN and PGE2. (One-way ANOVA, IL-6+PGE2 : IL-6+rmMTRN (1.8 mg/kg)+ PGE2, p-value = <0.0079, IL-6+PGE2 : Vehicle+PGE2, p-value = <0.0124) **E)** Grimacing behavior was observed to be similar to its baseline measurements in mice administered with rmMTRN (Two-way ANOVA, F = 10.35, p-value = <0.0001, Day1 post PGE2 injection, IL-6+PGE2 vs. Baseline measurement, p-value = 0.0302, Day2, IL-6+PGE2 vs. Baseline measurement, p-value = 0.0256, Day1 post PGE2 injection, IL-6+rmMTRN (1.8 mg/kg)+PGE2 vs. Baseline measurement of IL-6+ rmMTRN (1.8 mg/kg), p-value = 0.4465, Day2, IL-6+rmMTRN (1.8 mg/kg)+PGE2 vs. Baseline measurement of IL-6+rmMTRN (1.8 mg/kg), p-value = 0.9167).

## Discussion

The most important finding emerging from this work is that even after the resolution of overt pain-related behaviors caused by a treatment that causes neuropathic pain in mice and humans, changes in nociceptor gene expression at the level of mRNA translation persist. This finding suggests that these neurons do not readily return to their baseline state and that this altered translational landscape is linked to pain susceptibility where a normally innocuous stimulus can now precipitate a return of the pain state. We provide evidence in support of this hypothesis with GPR88, the most translationally upregulated mRNA in the DRG of primed mice, and with Meteorin, which was translationally downregulated. Using the IL-6 priming model, we demonstrate the increased translation of GPR88 causes IL-6-treated mice to become primed to a specific GPR88 agonist that is non-noxious in vehicle-treated mice. Conversely, enhancing Meteorin signaling with rmMTRN treatment reversed hyperalgesic priming. Collectively, our experiments demonstrate that persistent changes in nociceptor translation are a likely causative factor in the transition to a chronic pain state.

Translation regulation in nociceptors has long been recognized as a key mechanism that controls hyperalgesic priming (*13, 18–27, 62*). At least 3 translation regulation mechanisms have been proposed to control translation in nociceptors in priming models: MNK-eIF4E signaling (*12, 19, 29, 63*), the mechanistic target of rapamycin (mTOR) kinase signaling (*13, 46*), and translation regulation at the 3’ end of mRNAs through binding factors at the poly-A tail of mRNAs (*24, 27, 62*). While our study reveals mRNAs with altered translation in nociceptors in the primed state, we have not determined which of these pathways regulate the translation of specific subsets of mRNAs. Because these pathways may act in concert to control translation in sensitized nociceptors (*12*), it is also possible that interfering with these pathways individually would have similar consequences. We observed shifts in translation efficiency for roughly equal upregulated and downregulated mRNAs. This observation is consistent with MNK-eIF4E signaling which does not appear to decrease overall translation in mouse or human neurons but instead shifts subsets of mRNAs to increased translation efficiency and others to decreased efficiency (*56, 64–66*), as we observed in our study.

GPR88 is an orphan GPCR that does not have a known endogenous ligand or function, but several synthetic agonists have recently been described (*54*). The receptor has garnered interest in the neuropsychopharmacology area due to phenotypes of the *Gpr88* knockout mouse and genetic association studies that have implicated the receptor in psychiatric disorders (*54*). Interestingly from a pain perspective, GPR88 expression and activation seems to decrease the signaling of co-expressed mu-opioid receptors, including decreased supraspinal analgesia (*67*). Whether or not the receptor can also decrease signaling in peripheral opioid receptors has not been tested but would be of interest for future studies. While *Gpr88* mRNA is detected in mouse DRG in bulk RNA sequencing studies (*68, 69*), its expression level is very low and the cell types that express the mRNA are not clear from single nucleus sequencing experiments where it is mostly not detected (*11, 70*). It is also lowly expressed in human DRG (*68, 69, 71, 72*) but is detected in a subset of nociceptors that are associated with injury (*71*). Based on these findings, it is likely that the *Gpr88* mRNA is normally lowly expressed but its translation efficiency is increased after priming, leading to the increased association with translating ribosomes we detected in our study. The fact that a GPR88 agonist causes mechanical hypersensitivity and grimacing only in primed mice is consistent with this conclusion.

*Metrn* mRNA translation was decreased in primed DRG nociceptors, and systemic administration of Meteorin to mice led to reversal of hyperalgesic priming. Previous studies with Meteorin have suggested that this secreted protein with no known receptor alleviates pain in nerve injury models of neuropathic pain and in the paclitaxel model of chemotherapy-induced neuropathic pain in mice (*42, 60, 61*). This is the first study to demonstrate that Meteorin treatment can reverse hyperalgesic priming. Meteorin’s biological action has been linked to satellite glial cells and axonal network formation in previous studies, but the source of Meteorin protein has not been determined (*42, 59–61*). *Metrn* gene expression is detected in single nucleus RNA sequencing in *Scn10a* expressing cells in the mouse DRG (*70, 73*) and is also detected in these neurons in human DRG (*70, 71*). Our findings suggest that downregulation of the translation of the *Metrn* mRNA in nociceptors likely contributes to the primed state because treatment with Meteorin protein led to a complete and rapid reversal of hyperalgesic priming. This finding, combined with the previous literature on Meteorin in pain (*42, 60, 61*), suggests that Meteorin supplementation may be an effective strategy for pain treatment.

Our study has limitations. First, we focused entirely on female mice. We did this because paclitaxel is mostly used as a chemotherapeutic agent in women and because chronic pain is more prevalent in women (*74*). We acknowledge that future studies in male mice are warranted, but we also note that we have and others have already shown efficacy for Meteorin in both male and female animals suggesting no sex difference for this mediator in the context of pain (*42, 60, 61*). Second, our study focused on a limited number of genes, and a more comprehensive analysis of the entire translatome could uncover additional key players in the maintenance of chronic pain states. From this perspective, our work is a resource that others can use to explore these factors. Finally, we have only focused on nociceptors. It is known that immune cells play an important role in hyperalgesic priming and the promotion and resolution of chronic pain (*36–41, 75, 76*). No previous studies have used TRAP sequencing to identify persistent changes in translatomes of specific cells after pain has resolved from an initial stimulus. Our work sets a foundation for additional studies in this area of research.

The key takeaway from these experiments is that even after the resolution of overt signs of pain, nociceptors display a reorganization of their translational landscape that contributes to the maintenance of a primed state that makes these neurons susceptible to normally subthreshold stimuli that can now rekindle a state of pain in the organism driven by nociceptor hyperexcitability. These findings have important implications for understanding chronic pain states and for discovery of pain resolution mechanisms that can target the underlying pathology in nociceptors that maintain the susceptibility of the organism to experience pain in response to insults or stimuli that would normally not provoke such a response.

